# THz irradiation inhibits cell division by affecting actin dynamics

**DOI:** 10.1101/2021.02.26.433024

**Authors:** Shota Yamazaki, Yuya Ueno, Ryosuke Hosoki, Takanori Saito, Toshitaka Idehara, Yuusuke Yamaguchi, Chiko Otani, Yuichi Ogawa, Masahiko Harata, Hiromichi Hoshina

**Affiliations:** Terahertz Sensing and Imaging Research Team, RIKEN Center for Advanced Photonics, 519-1399 Aramaki-Aoba, Aoba-ku, Sendai, Miyagi 980-0845, Japan; Laboratory of Molecular Biology, Graduate School of Agricultural Science, Tohoku University, 468-1 Aramaki-Aoba, Aoba-ku, Sendai, Miyagi 980-0845, Japan; Research Center for Development of Far-Infrared Region, University of Fukui (FIR UF), Bunkyo 3-9-1, Fukui 910-8507, Japan; Laboratory of Bio-Sensing Engineering, Graduate School of Agriculture, Kyoto University, Kitashirakawa-Oiwakecho, Sakyo-ku, Kyoto 606-8502, Japan

## Abstract

Biological phenomena induced by terahertz (THz) irradiation are described in recent reports, but underlying mechanisms, structural and dynamical change of specific molecules are still unclear. In this paper, we performed time-lapse morphological analysis of human cells and found that THz irradiation halts cell division at cytokinesis. At the end of cytokinesis, the contractile ring, which consists of filamentous actin (F-actin), needs to disappear; however, it remained for 1 hour under THz irradiation. Induction of the functional structures of F-actin was also observed in interphase cells. Similar phenomena were also observed under chemical treatment (jasplakinolide), indicating that THz irradiation assists actin polymerization. We previously reported that THz irradiation enhances the polymerization of purified actin in vitro; our current work shows that it increases cytoplasmic F-actin in vivo. Thus, we identified one of the key biomechanisms affected by THz waves.

## Introduction

The recently developed technology of terahertz (THz) light sources indicate the bloom of applications in a wide range of fields, such as chemical sensing [1,2], security imaging motion sensing [3–6], and telecommunications [7–12]. For example, in the wireless technology “6G” aiming for practical use in the 2030s, the use of sub-THz electromagnetic waves is being studied. The use of the “sub-THz” is also being considered for the acquisition of high-precision position information in radars required for autonomous driving and motion sensors. Over the next decades, THz light sources will become miniaturized, powerful, cheap, and familiar to everyday life. To facilitate such practical THz applications, the safety of THz radiation for human health must be guaranteed [13].

The interaction between THz radiation and biological systems has been previously investigated. Two projects, the European THz-BRIDGE and the International EMF project in the SCENIHR [14], summarize recent studies about the biological effects of THz radiation. For example, THz irradiation was shown to inhibit cell proliferation and to change the adhesive properties of the nerve cell membrane [15,16]. Other studies showed THz-induced DNA destabilization [17–19], which causes chromosomal aberrations in human lymphocytes [20]. The transcriptional activation of wound-responsive genes in mouse skin [21] and the induction of DNA damage in an artificial human 3D skin tissue model [22] were also reported as effects of THz irradiation. However, the mechanisms are still unclear because such phenomenological studies cannot reveal the underlying molecular origin in the complex biological systems.

An important point to consider for THz irradiation experiments is the THz radiation source itself. The THz power density must not be too high to avoid detrimental thermal effects on the sample. Many studies have shown the effect of heating on cells, such as tissue damage [23,24], heat-induced cellular death [25,26], and DNA damage [27,28]. Thus, the THz beam should not be focused tightly to prevent an increase in the temperature on the sample. Two studies have shown that millimeter-wave radiation induces specific cellular responses that differ from direct thermal effects (29, 30); however, the underlying mechanism and exact targets are poorly defined. In addition to the effect of heating, the generation of the acoustic waves in aqueous solution must be considered when using the pulsed THz sources. In our previous works, we observed that THz pulses generate shockwaves at the surface of liquid water [31]. The generated shockwaves propagate to a depth of several millimeters, and disrupt protein structures in living cells [32]. To avoid such acoustic effects, the peak power of the THz pulses should be kept at a sufficiently low level.

In this study, we investigated the “non-thermal” and “non-acoustic” effects of THz irradiation on the morphology of living HeLa cells. The energy of THz was 6 mJ/cm^2^ with a duration of 10 ms, giving a peak power less than 0.6 W/cm^2^, which is eight orders of magnitude smaller than that in our previous studies [32]. The THz fluence was low enough to keep the temperature rise less than 0.2 °C during irradiation. Morphological observation showed that cell division in the cell cycle is arrested at mitosis during THz irradiation. Fluorescence microscopy revealed that this phenomenon is due to the stabilizing of the contractile ring, which is required to disappear to complete the cytokinesis — the last step of cell division. We found that the contractile ring was stabilized because of the enhancement of actin polymerization by THz irradiation. This work is the first to identify the key molecule and mechanism by which THz waves affect biological systems in a non-thermal and non-acoustic manner.

## Materials and Methods

### THz source

We used a gyrotron (FU CW GVIB [33]) to generate 0.46-THz waves. We designed an apparatus that exposed samples to the radiation, which had a peak power density of 0.6 W/cm^2^. A schematic representation of the device is shown in Fig. 1A. The THz gyrotron produced 10-ms-long pulses with a 1-Hz repetition rate [34]. As a second source of THz irradiation, we used a compact solid-state device based on an IMPATT-diode (TeraSense Group Inc), which ensured coherent continuous-wave emission of THz waves with a frequency of 0.28 THz and output power of 20 mW. THz radiation was outputted from the horn antenna (4 mm × 4 mm), and emitted from the bottom of the dish without focusing the beam, and with a power density of 125 mW/cm^2^.

**Fig. 1.**
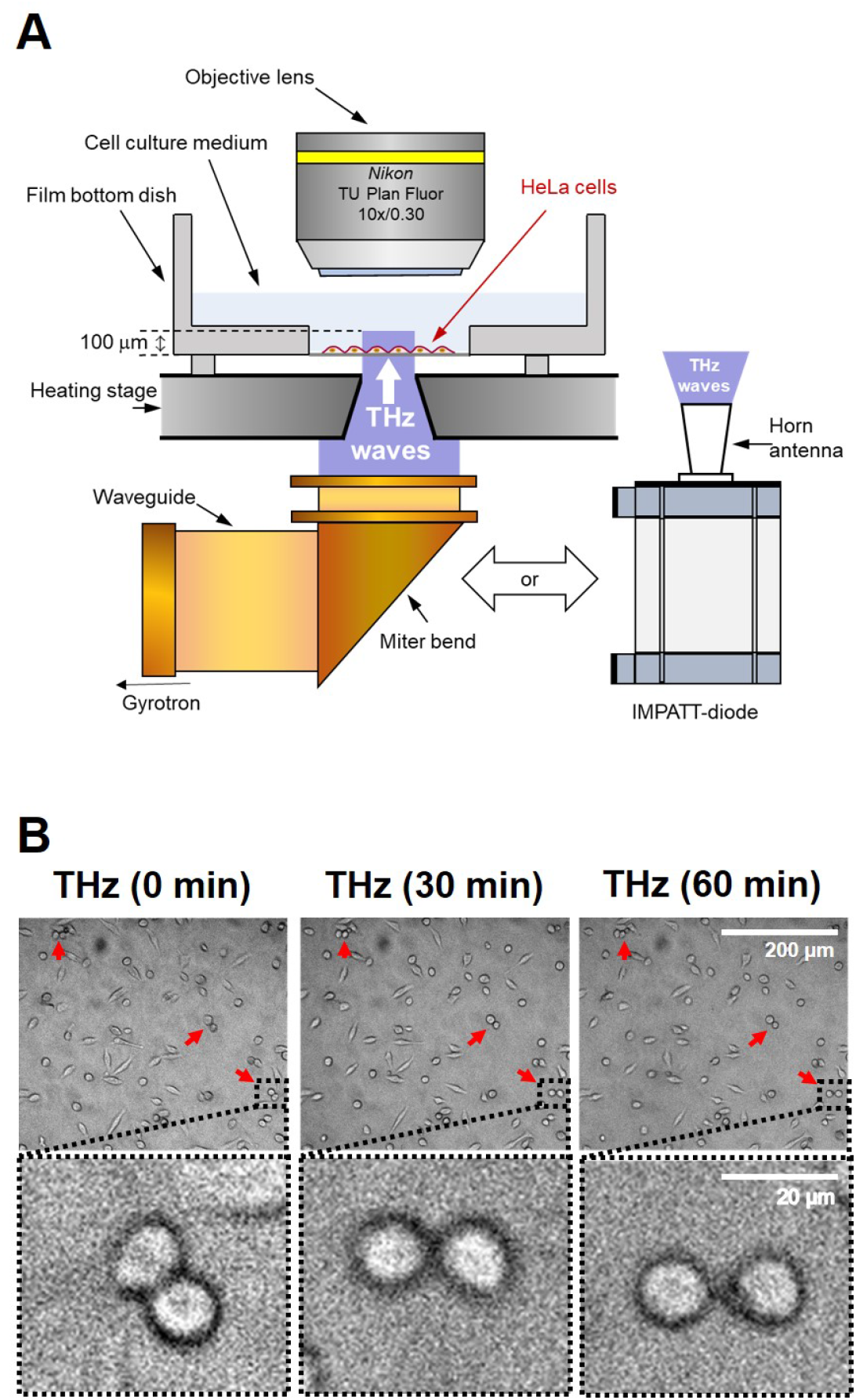
Effects of THz irradiation on cell morphology. (A) Schematic illustration of the experimental setup. THz waves with a power density of 0.6 W/cm^2^, frequency of 0.46 THz, pulse duration of 10 ms, and a repetition rate of 1 Hz were generated by a gyrotron at FIR-UF. The THz beam passed vertically from the bottom of the dish via an aperture of 4 mm in the heating stage. As a second source of THz irradiation, we used a IMPATT-diode which ensured coherent continuous-wave emission of THz waves with a frequency of 0.28 THz and output power of 20 mW. THz radiation was outputted from the horn antenna with a power density of 125 mW/cm^2^. HeLa cells were seeded on the film bottom dish and cultured for 24 hours before the experiments. The culture medium was kept at 37 °C by the heating stage during the experiments. (B) Microscopy images of cells at 0, 30, and 60 minutes. Irradiation was started at 0 minutes and continued for 60 minutes. The bottom panels show the magnified images of the black squares in the upper panels. The red arrows indicate a pair of cells with a round shape. The scale bar represents 200 μm (upper panel) and 20 μm (bottom panel).

### THz irradiation of HeLa cells

HeLa cells were seeded on 0.15 mm-thick cover glass and cultured in Dulbecco’s Modified Eagle’s Medium (Gibco) supplemented with 10% fetal bovine serum and antibiotics (penicillin and streptomycin) at 37 °C in a 5% CO_2_ humidified atmosphere. Actin filaments were stained with SiR-actin by adding probes from a 1 mM dimethyl sulfoxide (DMSO) stock solution to the growth medium (final concentration: 3 μM) and incubating for 1 hour at 37 °C in a 5% CO_2_ humidified atmosphere. The film dish was set on a heating stage (LINKAM: 10002L) to maintain a culture temperature at 37 ±1 °C. The THz beam passed vertically through a 4-mm hole in the heating stage. During THz irradiation, fluorescence microscopy images were obtained with a UV light source (Thorlabs, X-Cite 200DC lamp), dichroic mirror (Thorlabs, DMLP650R), two optical filters (excitation band pass: 625 nm/25 nm; emission long pass: 675 nm), objective lens (Olympus, LUMFLN60XW; Nikon, N10X-PF), and an sCMOS camera (Thorlabs, CS2100M-USB). Figure 1A shows a schematic diagram of the experimental setup for THz irradiation. Cells treated with 10 nM jasplakinolide in DMSO were used as a positive control.

For the quantitative analysis of the cells at cytokinesis, cells were synchronized at the mitotic phase using 25 μg/ml nocodazole. Cells were cultured at 16 hours after the addition of nocodazole. Before each experiment, nocodazole was washed out by changing the culture medium, and cells proceeded to mitosis with or without THz irradiation.

Image analysis was performed using Fiji software. To measure the mean signal intensity in the membrane compartment, the outline of each cell was selected using the area selection tools in the software. The mean signal intensity of the signal over the area of the cell was recorded. The number of cells is shown as *n*. Statistical significance was calculated using F- and T-tests.

### Morphological analysis

To measure the cell area and perimeter, the outlines of cells were selected (in the *x–y* plane) using the area selection tools in the Fiji software. The form factors of individual cells were calculated as 4*πS/L*^2^, where *S* is the projected cell area and *L* is the cell perimeter. This index reflects the irregularity of the cell shape: a perfectly round cell has a value of one, and a stellate cell has a value lower than one. Data are presented as the mean ± standard deviation. The number of cells is shown as *n*. Statistical significance was calculated using F- and T-tests.

## Results

### THz irradiation halts cell division of cultured cells

To observe the non-thermal and non-acoustic effects of the THz irradiation, we irradiated living cells with a THz beam with relatively low peak power. The sample was irradiated with the output of the gyrotron (0.46 THz), without focusing the beam and with a peak power density of 0.6 W/cm^2^. This radiation power is eight orders of magnitude lower than the power in which acoustic waves were generated in our previous work [32]. The radiation source was pulsed with a duty ratio of 1% (10-ms duration, 1-Hz repetition rate) to reduce heating of the sample. HeLa cells were grown on a film-bottom dish, and the culture medium was kept at 37 °C by a heating stage during the experiment. THz radiation was emitted from the bottom of the culture dish for 60 minutes (Fig. 1A). The high absorbance of water (160 cm^−1^ at 21 °C, 0.46 THz) limits the penetration depth of the THz waves to about 100 μm. Because the thickness of the cells is less than 30 μm, THz waves reached all regions of the cell. To evaluate the effect of THz irradiation, we performed time-lapse microscopy imaging of the HeLa cells (Fig. 1B).

Under THz irradiation, the appearance of a characteristic form of cells, which consists of a pair of round cells, was frequently observed (Fig. 1B, red arrow), and the characteristic cells are maintained during THz irradiation up to 60 min (Fig. 1B, bottom panel (zoomed images)). The round shape of the cells is a typical morphology of mitotic cells, and the pairing of two round cells is observed at the last step of mitosis, called cytokinesis (Fig. 2A). Cytokinesis is generally completed within 15 minutes [35]. Therefore, the persistence of the paired round cells indicates that THz irradiation inhibited the progression of cytokinesis.

**Fig. 2.**
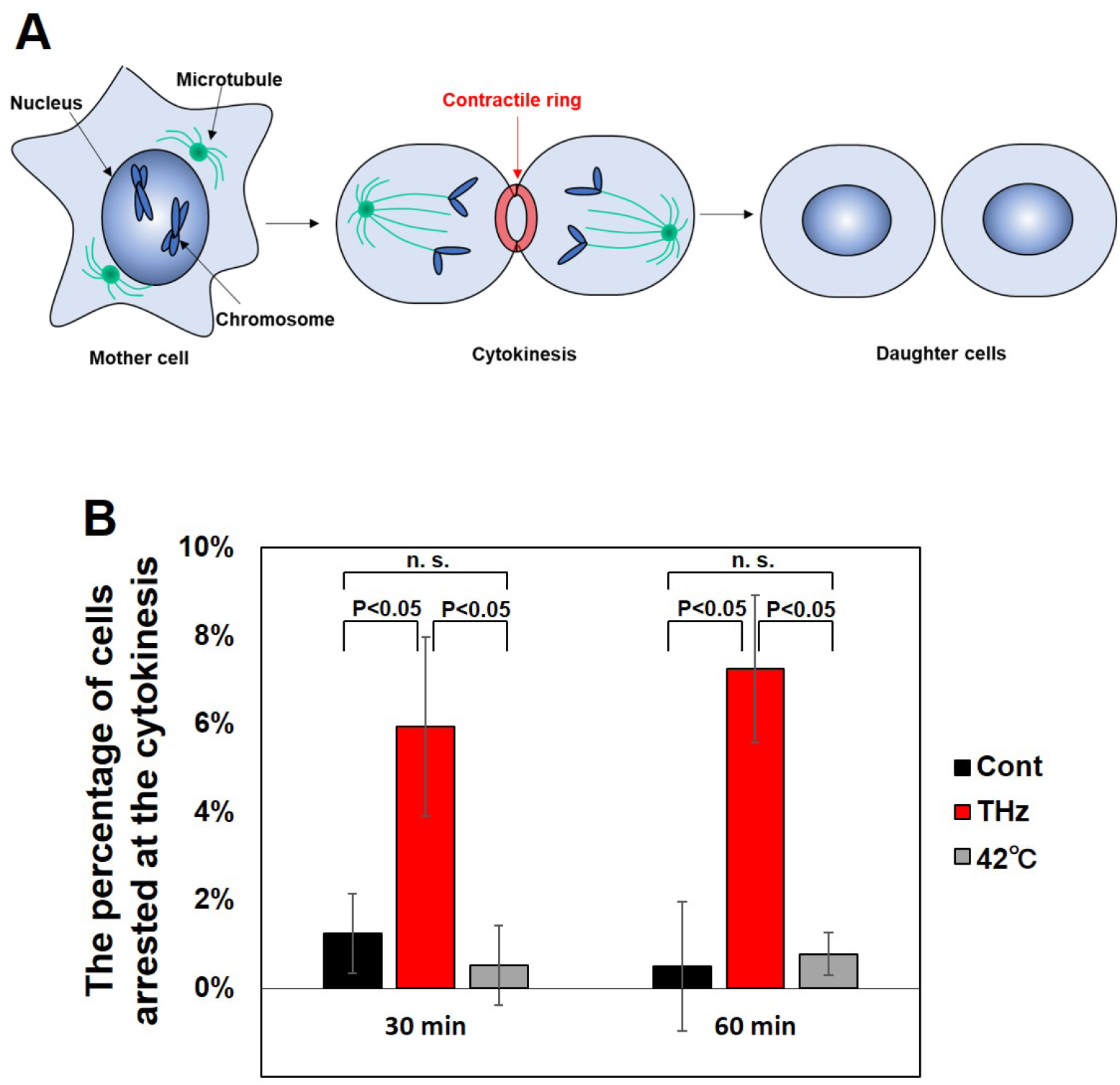
THz irradiation halts cytokinesis. (A) Schematic representation of mitotic progression. In the process of mitosis, actin polymerization is induced to make the contractile ring, which is required for starting the division of the mother cell into two daughter cells. Then, the contractile ring is squeezed and completes cell division. Cytokinesis is generally completed within 15 minutes. (B) Percentage of cells arrested at cytokinesis. The cell cycle was synchronized to the mitosis phase with 25 μg/ml nocodazole before each experiment. Nocodazole interferes with the polymerization of microtubules and arrests the initial step of mitosis. Cells were determined to be arrested at cytokinesis when the contractile ring was retained for more than 30 minutes after the release from nocodazole. The error bars show the standard deviation of three independent experiments. More than 184 cells were measured in each experiment.

For the quantitative evaluation of the arrested cells at cytokinesis, cells were synchronized at the initial phase of mitosis using 25 μg/ml nocodazole, and released into the culture medium without nocodazole to proceed with the mitosis. Figure 2B shows the percentage of cells arrested at cytokinesis. Whereas cytokinetic-arrested cells are not observed under the control condition, THz irradiation induced cytokinetic arrest at 30 minutes after nocodazole release and the arrest was further continued (Fig. 2B, THz). We also analyzed the effect of heat on the progression of cytokinesis. The culture medium was kept at 42 °C by the heating stage during the progression of mitosis; however, this did not increase the number of cells arrested at cytokinesis (Fig. 2B, 42 °C). Since the temperature rise during THz irradiation was less than 0.2 °C (Supplemental Fig. S1), some other reasons than the temperature increase are supposed for the inhibition of cytokinesis.

### Persistence of the contractile ring during THz irradiation

The dominant regulator of cytokinesis is the contractile ring, which consists of actin filaments (Fig. 2A) [36]. At the start of cytokinesis, a G (globular)- to F (filamentous)-actin transition is induced to make the contractile ring (polymerization reaction). Then, the opposite transition of F- to G-actin disassembles the contractile ring to complete cell division (depolymerization reaction). After THz irradiation, the percentage of cells arrested at cytokinesis significantly increased in comparison with control cells (Fig. 2B), suggesting that THz irradiation affects the disassembly of the contractile ring.

To observe the behavior of the contractile ring under THz irradiation, we stained actin filaments in living cells with SiR-actin [37], and performed time-lapse imaging under a fluorescence microscope. The formation of the contractile ring was observed with and without THz irradiation (Fig. 3, Cytokinesis, red arrow). Without THz irradiation, the contractile ring disappeared after 30 minutes, and the two daughter cells separated completely (Fig. 3, Control, white arrows). By contrast, under THz irradiation, the contractile ring remained for at least 30 minutes (Fig. 3, THz, 30 min later). In cells cultured at 42 °C, the contractile ring disappeared, and cell division was completed within 30 minutes (Fig. 3, 42 °C, 30 min later). This result suggests that the depolymerization reaction of actin progresses in a non-thermal manner. Cytokinesis is generally completed within 15 minutes, and the dynamic turnover of actin filaments to G-actin is required for its completion [38–41]. Importantly, the chemical induction of actin polymerization with jasplakinolide inhibits the completion of cytokinesis by stabilizing the contractile ring [42]. Taken together, these results support the notion that THz irradiation inhibits the completion of cytokinesis by affecting the actin dynamics.

**Fig. 3.**
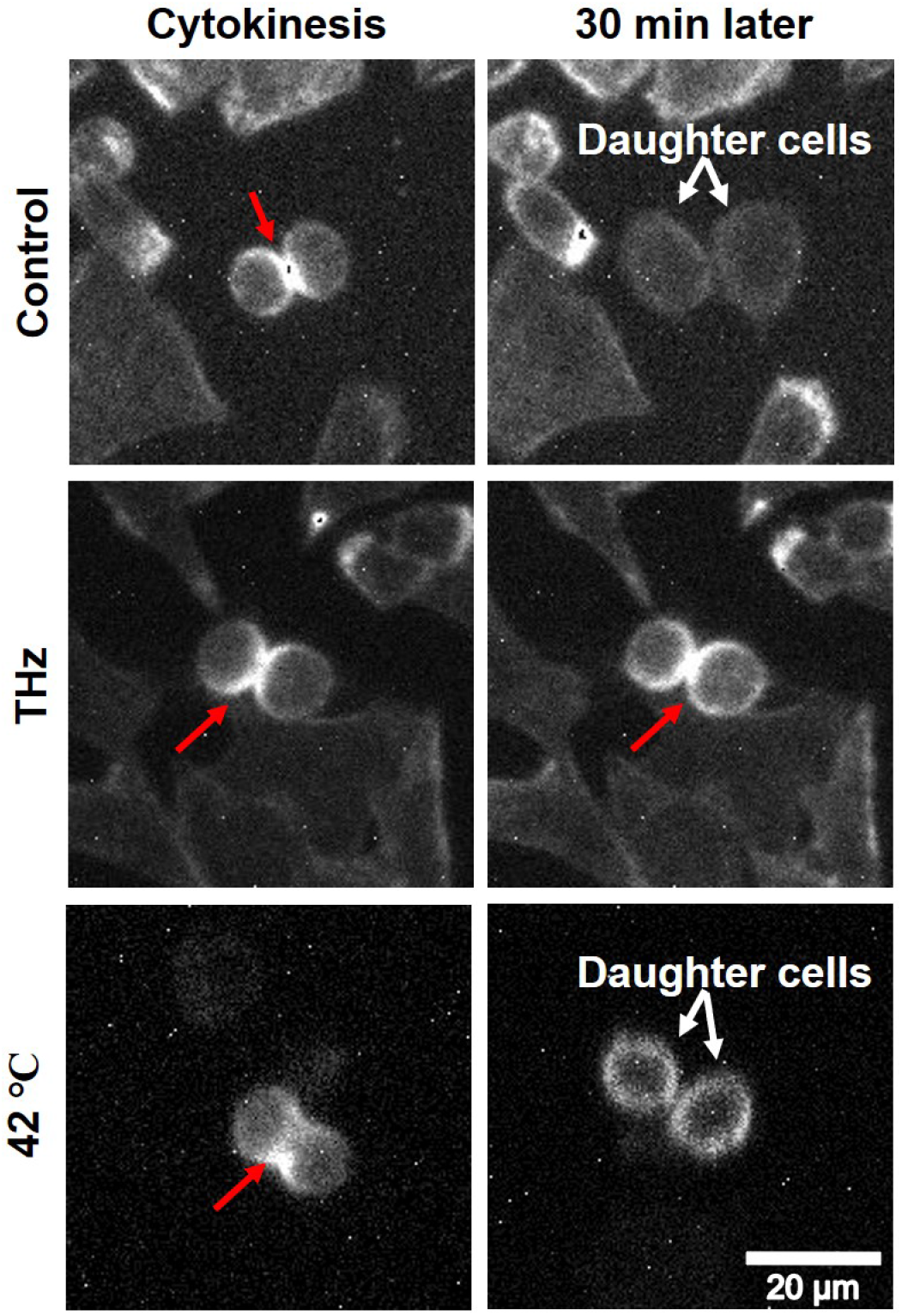
Persistence of the contractile ring during THz irradiation. Live-cell imaging of cells with a contractile ring. The cellular actin filaments were stained with SiR-actin and observed 30 minutes after generation of the contractile ring. The culture medium was kept at 37 °C by the heating stage during the experiments with and without THz irradiation. To observe the thermal effects on cytokinesis progression, cells were cultured at 42 °C and observed (bottom panel). The red arrows show the contractile ring and the white arrows show the daughter cells. The white bar shows a scale of 20 μm.

### Effects of THz irradiation on actin filaments inside cells

Actin filaments are relevant to various cellular functions, and their dynamics are tightly regulated. For example, cytoplasmic actin polymerization in the plasma membrane is an essential and versatile process that defines the cellular shape and confers mobility to cells. To evaluate the effects of THz irradiation on the actin dynamics observed in living cells, we stained actin filaments in living HeLa cells with SiR-actin [37], and performed time-lapse imaging with fluorescence microscopy. The fluctuation of the cellular actin filaments can be quantitatively estimated by the fluorescence intensity of SiR-actin, which increases up to 100-fold in the actin filaments. Cells treated with 10 nM jasplakinolide, which induce actin polymerization, were also analyzed as a positive control.

As Figure 4A shows, most cells stayed adherent during the 60-minute observation period, with a few cells detaching from the bottom of the dish. In addition, the area of the cells remained constant for 60 minutes during both THz irradiation and jasplakinolide treatment (Supplemental Fig. S2), suggesting that abnormal shape changes, such as atrophy and hypertrophy, did not occur. Figure 4B shows the mean fluorescence intensity of SiR-actin in individual cells at 0, 30, and 60 minutes. The box plot shows the mean fluorescence intensity of SiR-actin in the cells, and the error bar represents the standard deviation. The fluorescence intensities of SiR-actin in the cells were kept constant for 60 minutes in the control experiment (Fig. 4B, control), showing that fluorescence bleaching did not occur during the observation period. After 60 minutes of THz irradiation, the fluorescence intensity of SiR-actin increased, indicating that actin polymerization was accelerated and the number of filaments increased inside the cells (Fig. 4B, THz). A similar effect was observed for the ‘chemical’ induction of actin filamentation using jasplakinolide (Fig. 4B, Jasplakinolide). These results show that THz irradiation accelerates actin filamentation in living HeLa cells.

**Fig. 4.**
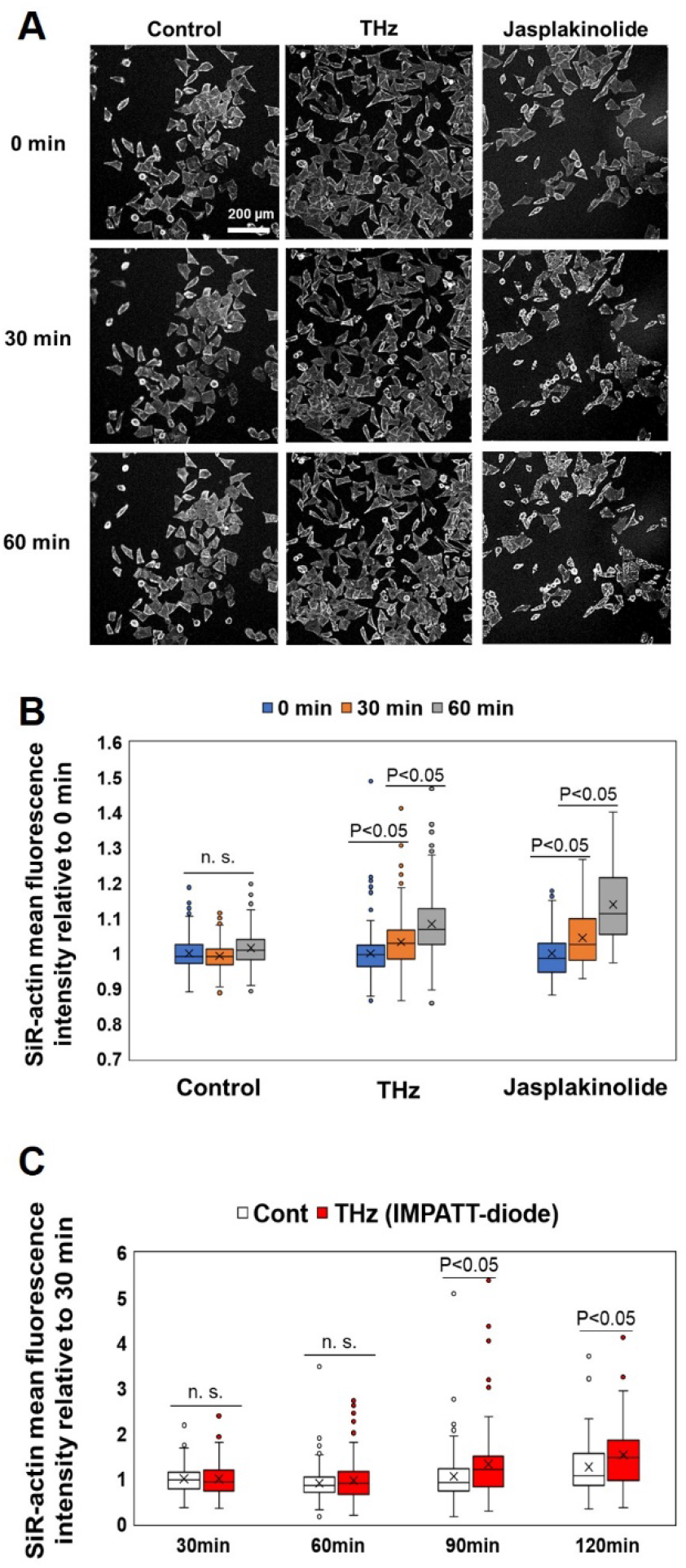
THz waves enhance actin polymerization in cells. (A) Fluorescence microscopy images of cells stained with SiR-actin at 0, 30, and 60 minutes. THz irradiation was started at 0 minutes and continued for 60 minutes. As a positive control, cells were treated with 10 nM jasplakinolide at 0 minutes to induce actin polymerization. The white bar shows a scale of 200 μm. (B) Mean fluorescence intensity of SiR-actin in individual cells measured from the fluorescence microscopy images. The box plot shows the mean value relative to 0 minutes. The standard deviations of three independent experiments are shown. More than 77 cells were measured in each experiment. (C) Irradiation with THz waves generated by the IMPATT-diode source was started at 0 minutes and continued for 120 minutes. The mean fluorescence intensity of SiR-actin in individual cells was measured from the fluorescence microscopy images. The box plot shows the mean value relative to that measured at 30 minutes. The standard deviations of three independent experiments are shown. More than 120 cells were measured in each experiment.

To confirm the THz irradiation effect by another type of radiation source, same experiment was performed by a solid-state semiconductor device (TeraSense: IMPATT diode), which outputs continuous-wave at 0.28 THz with a power of 20 mW. THz wave was emitted from the diagonal horn antenna with a size of 4 mm × 4 mm, attached at the bottom of the film-bottom dish (Fig. 1A). The irradiation power density was about 125 mW/cm^2^. Figure 4C shows the mean fluorescence intensity of SiR-actin in the individual cells at 30, 60, 90, and 120 minutes. After 90 minutes of irradiation, the fluorescence intensity of SiR-actin was significantly increased compared with the control cells (Fig. 4C, THz).

The fluorescence intensity under irradiation from the IMPATT diode increased more slowly than under gyrotron irradiation because of the different parameters of the two light sources. Specifically, the peak power of the IMPATT diode (125 mW/cm^2^) was about five times lower than that of the gyrotron (600 mW/cm^2^). Moreover, the frequency of the IMPATT diode (0.28 THz) was much lower than that of the gyrotron (0.46 THz). At present, we do not know which of these two parameters controls the speed of actin filamentation. We note that the average energy flux of the IMPATT diode (125 mJ/cm^2^/s) was higher than that of the gyrotron (6 mJ/cm^2^/s). However, the speed of actin filamentation does not depend on the average energy flux.

### Effects of THz irradiation on actin-including structures in interphase cells

In addition to the formation of the contractile ring in cytokinesis, actin polymerization is required for forming cellular structures in interphase cells, including stress fibers, lamellipodial meshworks, and transverse arcs (Fig. 5A). Stress fibers exist along the cell membrane and form the cytoskeleton, which maintains the cell shape. Lamellipodial meshworks are observed at the leading edge of cells and are required for cell migration. Transverse arcs are generated in the peripheral regions of the cell membrane and move to the center of the cell [43]; this movement is generally the initial step of cell migration, and actin polymerization is required for movement. To analyze the effect of THz irradiation on actin polymerization, we analyzed actin-including structures in living cells using fluorescent microscopy. Note that we did not observe any change of lamellipodial meshworks in this study. It is known that the production of lamellipodial meshworks induces the reorganization of the cell into an asymmetric shape. To confirm the cellular shape transition, we analyzed the form factor, which is close to 1 for a round shape, and close to 0 for an asymmetric shape [44]. The form factor was the same for the control, THz irradiation, and jasplakinolide-treated samples for 60 minutes (Supplemental Fig. S3). Therefore, we concluded that lamellipodial meshworks were not induced by the 60-minute THz irradiation.

**Figure 5.**
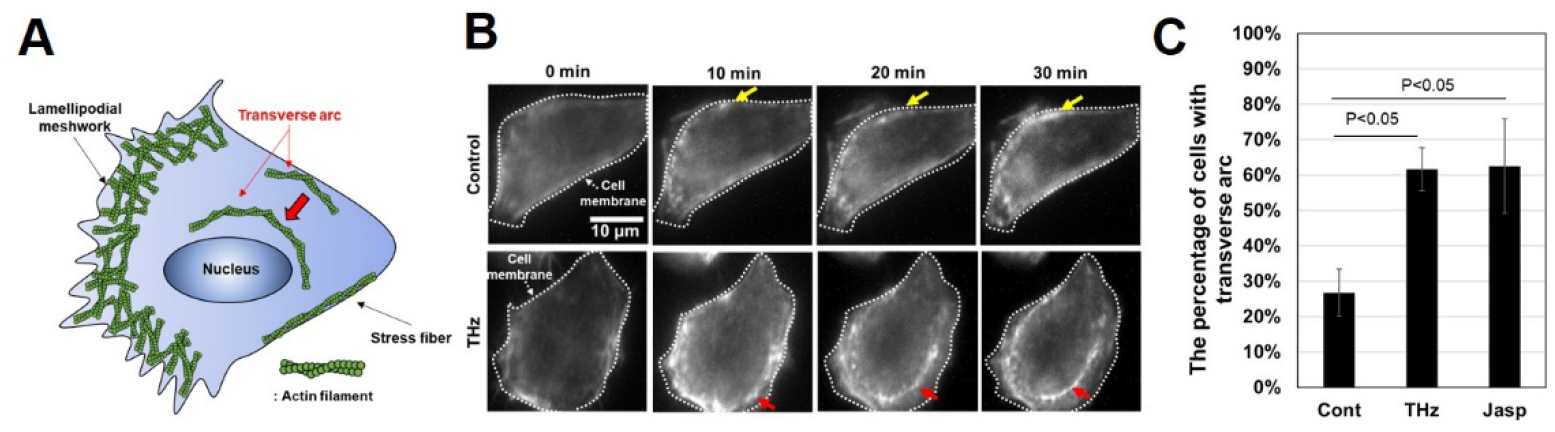
Effect of THz irradiation on actin-including structures. (A) Illustration of the functional structures that include actin filaments inside cells. In the cytoplasm, actin filaments form massive assemblies, which can be categorized as stress fibers, lamellipodial meshworks, and transverse arcs. Stress fibers are static structures that exist along the cell membrane; the lamellipodial meshwork is observed in the leading edge of the cell; and transverse arcs are generated in the cell membrane and move to the center of the cell. (B) Live-cell imaging of actin filaments with and without THz irradiation. The white dotted line marks the cell membrane. The yellow arrow shows stress fibers, which appeared along the cell membrane. The red arrow shows a transverse arc, which was generated in the cell membrane and moved to the center of the cell for 20–30 minutes. The scale bar represents 10 μm. (C) Percentage of cells, in which transverse arcs appeared during microscopy observation for 30 minutes. As a positive control, cells were treated with 10 nM jasplakinolide at 0 minutes to induce actin polymerization. The error bar shows the standard deviation of three independent experiments. More than 184 cells were measured in each experiment.

Figure 5B shows time-lapse images of a single cell stained with SiR-actin at 0, 10, 20, and 30 minutes. The white dotted lines show the position of the cell membrane. The fluorescence intensity of SiR-actin increased near the cell membrane in the control, indicating that stress fibers were generated during the measurement (Fig. 5B, Control, yellow arrows). Under THz irradiation, in addition to the stress fiber formation, transverse arcs were formed in the periphery, and this structure moves from the cell membrane towards the center of the cell (Fig. 5B, THz, red arrows) (Supplemental Movie. S1).

Figure 5C shows the number of cells in which transverse arcs were generated during the 30-minute experiment. 27% of cells contained a transverse arc in the control experiment (Fig. 5C, Cont). By contrast, over 60% of the cells contained a transverse arc as a result of either THz irradiation or jasplakinolide treatment (Fig. 5C, THz and Jasp). These results suggest that THz irradiation affects actin polymerization not only in the contractile ring but also in the cytoplasm of interphase cells.

## Discussion

In our previous study, we subjected an aqueous solution of purified actin protein to THz irradiation for the purpose of developing a physical technique for macromolecular manipulation [34]. In that study, we found that actin filaments were generated effectively under THz irradiation in living cells. Furthermore, THz irradiation caused the generation and retention of massive assemblies of actin filaments, such as contractile rings and transverse arcs (Figs 3 and 5).

Because the formation of biological molecules is sensitive to temperature, the simplest explanation for the enhancement of actin polymerization might be a transient increase of temperature owing to the absorption of THz irradiation by water. However, it has been demonstrated that the effect of a temperature rise on actin polymerization is negligible (Fig. 2B) [34,45]. In addition, we estimated the temperature change during THz irradiation as 0.23 °C using an adiabatic model (Supplemental Fig. S1). Therefore, it is unlikely that a temperature change due to THz irradiation enhances actin polymerization in living cells, and other mechanisms should be considered.

Another possible explanation is THz-induced shockwaves. In our previous study, we found that shockwaves were generated by THz pulses of 80 μJ/cm^2^ with a duration of 5 ps (peak power of 16 MW/cm^2^) [32]. Intense THz pulses are absorbed at the water surface and the energy concentration results in shockwave generation. The shockwaves propagate for a few millimeters in the aqueous medium, and disrupt the morphology of actin filaments in living cells. However, in the present study, the energy of each THz pulse was 6 mJ/cm^2^ with a duration of 10 ms, giving a peak power of just 0.6 W/cm^2^, which is eight orders of magnitude smaller than that used in Ref. 32, which generated shockwaves. Therefore, we consider that THz irradiation did not induce shockwaves under the experimental conditions of the present study.

We attribute the observed phenomena to non-thermal and non-acoustic effects of THz irradiation (i.e. the direct interaction between THz photons and the dynamical motion of the actin proteins). Because the vibration frequencies of the higher-order conformations of proteins and the surrounding water molecules are in the THz band [46–48], THz irradiation perturbates the intra- and inter-molecular dynamics of the actin proteins. The actin polymerization process consists of three phases: nucleation, elongation, and equilibrium. In our previous study, we found that THz irradiation enhances actin polymerization reaction in the aqueous solution [34]. We concluded that THz irradiation accelerates the elongation process because the actin filaments undergo additional elongation under THz irradiation in the equilibrium state. Those results showed that THz irradiation affects the dynamics of actin molecules during the elongation reaction.

Our previous in vitro THz irradiation experiment for the same molecule helps us understand the mechanism of in vivo THz irradiation. The observed phenomena — the inhibition of cytokinesis and formation of transverse arcs — suggest the enhancement of actin filamentation in living cells, which we also quantitatively confirmed from the fluorescence intensity of SiR-actin. In the in vitro experiment, such enhancement of actin filamentation was not due to the expression of the intra-cellular system, such as activation of cell signaling, changes of transcriptional regulations, and induction of cellular responses, but was due to the direct enhancement of the elongation reaction of the actin filament. Using actin molecules, we succeeded in elucidating the effects of THz irradiation on molecular reactions and cellular expression.

Actin filament is a major component of the cytoskeleton, and has crucial roles in determining cell shape, and for cell motility and division [49,50]. Moreover, the recent development of fluorescence probes has led to the revelation that nuclear actin filaments are required for transcriptional regulation, DNA repair, and gene reprogramming [51–53]. Therefore, THz irradiation has potential as a novel biological tool. In fact, we discovered that the effect of THz irradiation is similar to that of jasplakinolide treatment. Jasplakinolide, a naturally occurring cyclic peptide from the marine sponge Jaspis sp [54], is a membrane-permeable, actin-polymerizing, and filament-stabilizing drug [55]. Jasplakinolide has a wide range of known biological functions, which include antifungal and antitumor activities [56–58]. Thus, by analogy with jasplakinolide, we suggest that THz irradiation can be used to manipulate cell functions via actin polymerization. In this study, we also demonstrated that the actin filamentation is induced by an IMPATT diode source. The IMPATT diode is small, operated at room temperature, and works with lower electrical power. Such solid-state semiconductor THz-sources are widely available for experiments with biological samples.

## Conclusions

We found that THz irradiation enhances the formation and stabilization of actin assemblies in living cells. Therefore, we propose that THz irradiation can be used for the optical manipulation of cellular functions via the modulation of actin dynamics, leading to a better understanding of the function of actin.

## Acknowledgment

We thank Adam Brotchie, PhD, from Edanz Group (https://en-author-services.edanz.com/ac) for editing a draft of this manuscript and helping to draft the abstract.

## Supporting information

**S1 Fig. Temperature change of the sample due to THz irradiation.**

**S2 Fig. Morphological analysis of cells.**

**S3 Fig. Morphological analysis of cells.**

**S1 Movie. Live-cell imaging of actin filaments with THz irradiation.**

## References

1. Peiponen K-E, Zeitler A, Kuwata-Gonokami M. Terahertz spectroscopy and imaging. Springer; 2012.

2. Ueno Y, Ajito K. Analytical Terahertz Spectroscopy. Anal Sci. 2008;24: 185–192. doi:10.2116/analsci.24.185

3. Mittleman DM. Perspective: Terahertz science and technology. J Appl Phys. 2017;122: 230901. doi:10.1063/1.5007683

4. Dean P, Valavanis A, Keeley J, Bertling K, Lim YL, Alhathlool R, et al. Terahertz imaging using quantum cascade lasers—a review of systems and applications. J Phys D Appl Phys. 2014;47: 374008. doi:10.1088/0022-3727/47/37/374008

5. Guillet JP, Recur B, Frederique L, Bousquet B, Canioni L, Manek-Hönninger I, et al. Review of terahertz tomography techniques. J Infrared, Millimeter, Terahertz Waves. 2014;35: 382–411. doi:10.1007/s10762-014-0057-0

6. Lien J, Gillian N, Karagozler ME, Amihood P, Schwesig C, Olson E, et al. Soli: Ubiquitous gesture sensing with millimeter wave radar. ACM Trans Graph. 2016;35: 1–19. doi:10.1145/2897824.2925953

7. Giordani M, Polese M, Mezzavilla M, Rangan S, Zorzi M. Toward 6G Networks: Use Cases and Technologies. IEEE Commun Mag. 2020;58: 55–61. doi:10.1109/MCOM.001.1900411

8. Mourad A, Yang R, Lehne PH, De La Oliva A. A baseline roadmap for advanced wireless research beyond 5G. Electron. 2020;9: 1–14. doi:10.3390/electronics9020351

9. Saad W, Bennis M, Chen M. A Vision of 6G Wireless Systems: Applications, Trends, Technologies, and Open Research Problems. IEEE Netw. 2020;34: 134–142. doi:10.1109/MNET.001.1900287

10. Nagatsuma T. Advances in Terahertz Communications Accelerated by Photonics Technologies. OECC/PSC 2019 - 24th Optoelectron Commun Conf Conf Photonics Switch Comput 2019. 2019; 1–3. doi:10.23919/PS.2019.8818026

11. Koenig S, Lopez-Diaz D, Antes J, Boes F, Henneberger R, Leuther A, et al. Wireless sub-THz communication system with high data rate. Nat Photonics. 2013;7: 977–981. doi:10.1038/nphoton.2013.275

12. O’Hara JF, Ekin S, Choi W, Song I. A Perspective on Terahertz Next-Generation Wireless Communications. Technologies. 2019;7: 43. doi:10.3390/technologies7020043

13. Jamshed MA, Heliot F, Brown TWC. A Survey on Electromagnetic Risk Assessment and Evaluation Mechanism for Future Wireless Communication Systems. IEEE J Electromagn RF Microwaves Med Biol. 2020;4: 24–36. doi:10.1109/JERM.2019.2917766

14. Leitgeb N, Auvinen A, Danker-hopfe H, Mild KH. SCENIHR (Scientific Committee on Emerging and Newly Identified Health Risks), Potential health effects of exposure to electromagnetic fields (EMF), Scientific Committee on Emerging and Newly Identified Health Risks SCENIHR Opinion on Potential health. 2016. doi:10.2772/75635

15. Munzarova AF, Kozlov AS, Zelentsov EL. Effect of terahertz laser irradiation on red blood cells aggregation in healthy blood. Vestn NSU Phys Ser. 2013;8: 117–123.

16. Olshevskaya JS, Kozlov AS, Petrov AK, Zapara TA, Ratushnyak AS. Cell membrane permeability under the influence of terahertz (submillimeter) laser radiation. Vestn Novosib State Univ. 2010;5: 177–181.

17. Bock J, Fukuyo Y, Kang S, Lisa Phipps M, Alexandrov LB, Rasmussen KO, et al. Mammalian stem cells reprogramming in response to terahertz radiation. PLoS One. 2010;5: 8–13. doi:10.1371/journal.pone.0015806

18. Alexandrov BS, Rasmussen KØ, Bishop AR, Usheva A, Alexandrov LB, Chong S, et al. Non-thermal effects of terahertz radiation on gene expression in mouse stem cells. Biomed Opt Express. 2011;2: 2679. doi:10.1364/BOE.2.002679

19. Alexandrov BS, Lisa Phipps M, Alexandrov LB, Booshehri LG, Erat A, Zabolotny J, et al. Specificity and Heterogeneity of Terahertz Radiation Effect on Gene Expression in Mouse Mesenchymal Stem Cells. Sci Rep. 2013;3: 1–8. doi:10.1038/srep01184

20. Korenstein-Ilan A, Barbul A, Hasin P, Eliran A, Gover A, Korenstein R. Terahertz Radiation Increases Genomic Instability in Human Lymphocytes. Radiat Res. 2008;170: 224–234. doi:10.1667/RR0944.1

21. Kim KT, Park J, Jo SJ, Jung S, Kwon OS, Gallerano GP, et al. High-power femtosecond-terahertz pulse induces a wound response in mouse skin. Sci Rep. 2013;3: 1–7. doi:10.1038/srep02296

22. Titova L V., Ayesheshim AK, Golubov A, Fogen D, Rodriguez-Juarez R, Hegmann FA, et al. Intense THz pulses cause H2AX phosphorylation and activate DNA damage response in human skin tissue. Biomed Opt Express. 2013;4: 559. doi:10.1364/BOE.4.000559

23. Henriques FC, Moritz AR. Studies of Thermal Injury: I. The Conduction of Heat to and through Skin and the Temperatures Attained Therein. A Theoretical and an Experimental Investigation. Am J Pathol. 1947;23: 530–49. Available: http://www.ncbi.nlm.nih.gov/pubmed/19970945

24. Pearce JA. Models for Thermal Damage in Tissues: Processes and Applications. Crit Rev Biomed Eng. 2010;38: 1–20. doi:10.1615/CritRevBiomedEng.v38.i1.20

25. Boreham DR, Mitchel REJM. Heat-induced thermal tolerance and radiation resistance to apoptosis in human lymphocytes. Biochem Cell Biol. 1997;75: 393–397. doi:10.1139/o97-077

26. Wilmink GJ, Opalenik SR, Beckham JT, Abraham AA, Nanney LB, Mahadevan-Jansen A, et al. Molecular Imaging-Assisted Optimization of Hsp70 Expression during Laser-Induced Thermal Preconditioning for Wound Repair Enhancement. J Invest Dermatol. 2009;129: 205–216. doi:10.1038/jid.2008.175

27. Mori E, Takahashi A, Ohnishi T. The Biology of Heat-induced DNA Double-Strand Breaks. Therm Med. 2008;24: 39–50. doi:10.3191/thermalmed.24.39

28. Roti Roti JL, Pandita RK, Mueller JD, Novak P, Moros EG, Laszlo A. Severe, short-duration (0–3 min) heat shocks (50–52°C) inhibit the repair of DNA damage. Int J Hyperth. 2010;26: 67–78. doi:10.3109/02656730903417947

29. Habauzit D, Le Quément C, Zhadobov M, Martin C, Aubry M, Sauleau R, et al. Transcriptome Analysis Reveals the Contribution of Thermal and the Specific Effects in Cellular Response to Millimeter Wave Exposure. Scarfi MR, editor. PLoS One. 2014;9: e109435. doi:10.1371/journal.pone.0109435

30. Romanenko S, Harvey AR, Hool L, Fan S, Wallace VP. Millimeter Wave Radiation Activates Leech Nociceptors via TRPV1-Like Receptor Sensitization. Biophys J. 2019;116: 2331–2345. doi:10.1016/j.bpj.2019.04.021

31. Tsubouchi M, Hoshina H, Nagai M, Isoyama G. Plane photoacoustic wave generation in liquid water using irradiation of terahertz pulses. Sci Rep. 2020;10: 18537. doi:10.1038/s41598-020-75337-6

32. Yamazaki S, Harata M, Ueno Y, Tsubouchi M, Konagaya K, Ogawa Y, et al. Propagation of THz irradiation energy through aqueous layers: Demolition of actin filaments in living cells. Sci Rep. 2020;10: 1–10. doi:10.1038/s41598-020-65955-5

33. Idehara T, Tatematsu Y, Yamaguchi Y, Khutoryan EM, Kuleshov AN, Ueda K, et al. The development of 460 GHz gyrotrons for 700 MHz DNP-NMR spectroscopy. J Infrared, Millimeter, Terahertz Waves. 2015;36: 613–627.

34. Yamazaki S, Harata M, Idehara T, Konagaya K, Yokoyama G, Hoshina H, et al. Actin polymerization is activated by terahertz irradiation. Sci Rep. 2018;8. doi:10.1038/s41598-018-28245-9

35. Milo R, Phillips R. Cell Biology by the Numbers. Garland Science; 2015. doi:10.1201/9780429258770

36. Bezanilla M, Gladfelter AS, Kovar DR, Lee WL. Cytoskeletal dynamics: A view from the membrane. J Cell Biol. 2015;209: 329–337. doi:10.1083/jcb.201502062

37. D’Este E, Hell SW, Waldmann H, Göttfert F, Gerlich DW, Masharina A, et al. Fluorogenic probes for live-cell imaging of the cytoskeleton. Nat Methods. 2014;11: 731–733. doi:10.1038/nmeth.2972

38. Pelham RJ, Chang F. Actin dynamics in the contractile ring during cytokinesis in fission yeast. Nature. 2002;419: 82–86. doi:10.1038/nature00999

39. Group CB. Single Particle Tracking of Surface Receptor. 1994;127: 963–971.

40. O’Connell CB, Warner AK, Wang Y li. Distinct roles of the equatorial and polar cortices in the cleavage of adherent cells. Curr Biol. 2001;11: 702–707. doi:10.1016/S0960-9822(01)00181-6

41. Murthy K, Wadsworth P. Myosin-II-Dependent Localization and Dynamics of F-Actin during Cytokinesis. 2005;15: 724–731. doi:10.1016/j.cub.2005.02.055

42. Mendes Pinto I, Rubinstein B, Kucharavy A, Unruh JR, Li R. Actin Depolymerization Drives Actomyosin Ring Contraction during Budding Yeast Cytokinesis. Dev Cell. 2012;22: 1247–1260. doi:10.1016/j.devcel.2012.04.015

43. Vallenius T. Actin stress fibre subtypes in mesenchymal-migrating cells. Open Biol. 2013;3. doi:10.1098/rsob.130001

44. Altankov G, Grinneu F. Depletion of Intracellular Potassium Disrupts Coated Pits and Reversibly Inhibits Cell Polarization During Fibroblast Spreading. 1993;120.

45. Kawamura M, Maruyama K. A Further Study of Electron Microscopic Particle Length of F-Actin Polymerized *in Vitro*. J Biochem. 1972;72: 179–188.

46. Yamamoto N, Ohta K, Tamura A, Tominaga K. Broadband Dielectric Spectroscopy on Lysozyme in the Sub-Gigahertz to Terahertz Frequency Regions: Effects of Hydration and Thermal Excitation. J Phys Chem B. 2016;120: 4743–4755. doi:10.1021/acs.jpcb.6b01491

47. Xu Y, Havenith M. Perspective: Watching low-frequency vibrations of water in biomolecular recognition by THz spectroscopy. J Chem Phys. 2015;143: 170901. doi:10.1063/1.4934504

48. Conti Nibali V, Havenith M. New Insights into the Role of Water in Biological Function: Studying Solvated Biomolecules Using Terahertz Absorption Spectroscopy in Conjunction with Molecular Dynamics Simulations. J Am Chem Soc. 2014;136: 12800–12807. doi:10.1021/ja504441h

49. Pollard TD, Cooper JA. Actin, a central player in cell shape and movement. Science. 2009;326: 1208–1212. doi:10.1126/science.1175862

50. Small JV, Rottner K, Kaverina I, Anderson KI. Assembling an actin cytoskeleton for cell attachment and movement. Biochim Biophys Acta - Mol Cell Res. 1998;1404: 271–281. doi:10.1016/S0167-4889(98)00080-9

51. Yamazaki S, Yamamoto K, de Lanerolle P, Harata M. Nuclear F-actin enhances the transcriptional activity of β-catenin by increasing its nuclear localization and binding to chromatin. Histochem Cell Biol. 2016;145: 389–399. doi:10.1007/s00418-016-1416-9

52. Caridi CP, Plessner M, Grosse R, Chiolo I. Nuclear actin filaments in DNA repair dynamics. Nat Cell Biol. 2019;21: 1068–1077. doi:10.1038/s41556-019-0379-1

53. Miyamoto K, Pasque V, Jullien J, Gurdon JB. Nuclear actin polymerization is required for transcriptional reprogramming of Oct4 by oocytes. Genes Dev. 2011;25: 946–958. doi:10.1101/gad.615211

54. White KN, Tenney K, Crews P. The Bengamides: A Mini-Review of Natural Sources, Analogues, Biological Properties, Biosynthetic Origins, and Future Prospects. J Nat Prod. 2017;80: 740–755. doi:10.1021/acs.jnatprod.6b00970

55. Bubb MR, Senderowicz AMJ, Sausville EA, Duncan KLK, Korn ED. Jasplakinolide, a cytotoxic natural product, induces actin polymerization and competitively inhibits the binding of phalloidin to F-actin. J Biol Chem. 1994;269: 14869–14871.

56. Scott VR, Boehme R, Matthews TR. New class of antifungal agents: Jasplakinolide, a cyclodepsipeptide from the marine sponge, Jaspis species. Antimicrob Agents Chemother. 1988;32: 1154–1157. doi:10.1128/AAC.32.8.1154

57. Stingl J, Andersen RJ, Emerman JT. In vitro screening of crude extracts and pure metabolites obtained from marine invertebrates for the treatment of breast cancer. Cancer Chemother Pharmacol. 1992;30: 401– 406. doi:10.1007/BF00689969

58. Senderowicz AM, Kaur G, Sainz E, Laing C, Inman WD, Rodríguez J, et al. Jasplakinolide’s inhibition of the growth of prostate carcinoma cells in vitro with disruption of the actin cytoskeleton. J Natl Cancer Inst. 1995;87: 46–51. doi:10.1093/jnci/87.1.46

